# Does female post-copulatory preference depend on pre-copulatory choice and post-mating environment?

**DOI:** 10.1101/2025.03.10.642410

**Authors:** Krish Sanghvi, Biliana Todorova, Irem Sepil

## Abstract

Sexual selection operates across pre- and post-copulatory episodes, driven by intra-sexual competition and inter-sexual choice for mates or gametes. In females, sexual selection often manifests as choosiness, however pre- and post-copulatory preferences can be in opposing directions. While relationships between male pre- and post-copulatory traits are well-studied, these relationships are less understood in females. Additionally, female post-copulatory environments can potentially influence post-copulatory choosiness, but this has been little investigated. Using *Drosophila melanogaster*, we explored whether mating latency, a proxy for inter-sexual pre-copulatory choosiness, correlates with female ejaculate ejection behaviour, a proxy for post-copulatory choosiness. We further tested whether the presence of a male in the female’s post-copulatory environment influences her ejection behaviour. We found no significant effect of male presence. However, males with longer mating latencies experience a smaller proportion of their sperm ejected, suggesting that males preferred as mates may produce sperm less favoured for fertilization. This finding might possible trade-offs between male investment in courtship and ejaculates. Our study highlights that female-mediated sexual selection at pre- and post-copulatory stages can shape sexual traits in complex ways. This has implications for sexual conflict, possibly providing an explanation for the maintenance of variation in sexually selected traits.

## Introduction

Sexual selection occurs at both pre- and post-copulatory stages, each partitioned into intra-sexual competition between, and inter-sexual preference for, mates or gametes respectively (Kuijper et al, 2012). Females often mediate sexual selection both before and after copulation via preference for specific mating partners or their sperm. In females, pre-copulatory choosiness is typically expressed through behaviours such as preferring to mate with males with higher quality phenotypes (Dougherty, 2020; Edward, 2014; Edward and Chapman, 2011) or accepting matings from preferred males more quickly than from less preferred males (Davies et al, 2020; Reinhold et al, 2015). Post-copulatory choosiness (“cryptic female choice”) further allows females to bias paternity towards preferred males by modulating sperm use (Eberhard, 2015; Firman et al, 2017). For instance, females in some species eject sperm sooner or eject more sperm from a less-preferred male to reduce his paternity share (Pizzari et al, 2000). Both pre- and post-copulatory mechanisms act together to shape male and female sexually selected behaviours and traits.

Episodes of pre- and post-copulatory sexual selection can act synergistically or disruptively (Devigili et al, 2015; Marie-Orleach et al, 2024; McDonald et al, 2017; Sanghvi et al, 2024). For example, a male preferred by females during pre-copulatory inter-sexual mate choice (e.g. due to better courtship ability, more ornamentation, higher condition), might also be preferred post-copulation, leading to females retaining more sperm from these males for fertilisation (e.g. Parker, 2009; Sbilordo and Martin, 2014). Conversely, females coerced into mating by less-preferred males might eject their sperm sooner or eject more of their sperm (Dean et al, 2011) to prevent fertilisation. Furthermore, trade-offs between male investment in pre-copulatory traits that increase mating success, versus post-copulatory traits that increase fertilisation success, could lead to negative correlations between the direction of pre- and post-copulatory episodes of sexual selection (Danielsson, 2001; Evans and Garcia-Gonzalez, 2016; Ferrandiz-Roviro et al, 2014; Kim and Velando, 2020). Understanding how both, pre- and post-copulatory sexual selection influence reproductive success requires examining the relationship between pre-copulatory traits and post-copulatory behaviours (Kvanermo and Simmons, 2013). Some studies have assessed these relationships on traits mostly mediated by males, such as male weaponry and sperm competitiveness or testes size (e.g. Durrant et al, 2016; Locatello et al, 2006; Puniamoorthy et al, 2012). However, a studies investigating female pre- and post-copulatory inter-sexual choosiness are lacking (however, see Pilastro et al, 2004).

The fruit fly, *Drosophila melanogaster*, provides an excellent model to study pre- and post-copulatory sexual selection. Male fruit flies perform elaborate courtship displays that females assess before mating (Pan et al, 2011; Pavlou and Goodwin, 2013). Mating latency is often used as a proxy for female preference/choosiness, and males in higher condition or better at courtship typically have shorter mating latencies (e.g. Dukas, 2005; Hosken et al, 2008; Poissonnier et al, 2024; Savic-Veselinovic et al, 2017; Sharma et al, 2010). Post-copulation, females store sperm in the seminal receptacle and spermathecae, and eject the un-stored sperm along with the male’s mating plug within a few hours (Manier et al, 2010; Schnackenberg et al, 2012). This mating plug temporarily prevents female re-mating, thereby reducing sperm competition (Avila et al, 2015; Bretman et al, 2020; McDonough-Goldstein et al, 2022). Females can control the timing of ejaculate ejection (Laturney and Billeter, 2016; Lupold et al, 2013; Mahdjoub et al, 2023; Schnackenberg et al, 2012), based on their socio-sexual environment and the mated male’s quality. For instance, females eject mating plugs sooner when they can smell the cuticular pheromones of more attractive males, compared to pheromones of less attractive males, or when these pheromones are absent (Doubovetzky et al, 2024; Yun et al, 2024). While female ejection latency is assumed to be correlated with numbers of sperm stored (Doubovetzky et al, 2024; Manier et al, 2020), direct tests for this association are lacking. Additionally, whether the female’s post-mating environment (e.g. the presence of males) might modulate the relationship between pre- and post-copulatory traits as well as female post-mating behaviour remains unknown.

Using *D. melanogaster,* we address three aims. First, we test how pre-copulatory female choosiness, measured as mating latency, correlates with post-copulatory female behaviours. Second, we investigate how the presence of another male in the female’s post-mating environment influences her ejaculate ejection behaviour. Third, we quantify whether different measurements of female post-copulatory behaviour (i.e. ejection latency, sperm numbers stored, sperm numbers ejected, and sperm proportion ejected) correlate with each other. For aim 1, we predict that if pre- and post-copulatory selection act complementarily, such that males preferred before mating are also preferred after mating, then males with shorter mating latencies will also have more of their sperm retained for fertilization, and their mating plugs ejected later. Alternatively, if there are trade-offs between male investment in courtship versus ejaculates (Simmons et al, 2017), then males with shorter mating latencies will have their mating plugs removed earlier, have more sperm ejected, or have fewer sperm retained by the female. For aim 2, we predict that females encountering a second male after her first mating will eject the first male’s ejaculate sooner and in greater amounts if: (i) doing so allows her to remate sooner, increasing the genetic diversity of her offspring (Jennions and Petrie, 2000); or (ii) the presence of a second male reduces the risk of sperm depletion compared to when no additional mates are available. This is based on fruit fly females often facing a high risk of sperm depletion due to the low sperm-to-egg ratio (∼30:1-Bjork and Pitnick, 2006) and males mating polygynously (Sanghvi et al, 2025). Alternatively, females encountering another male may delay ejaculate ejection and retain more sperm if male harassment limits their ability to remove the mating plug. For aim 3, we predict that earlier ejection of ejaculates should lead to fewer sperm being stored, therefore more sperm being ejected by females, and a negative correlation between sperm stored versus sperm ejected. However, if males vary in the total quantities of ejaculate transferred to females, then males who transfer more sperm would not only have greater numbers of sperm stored by females, but also more sperm ejected by females. This would lead to a positive or a lack of correlation between sperm stored and ejected, following the logic of condition dependence versus trade-offs in life-history traits (Roff and Fairbairn, 2007).

## Methods

### Stock maintenance

We used *D. melanogaster* populations from three different lines for our experiment: Dahomey (*dah*), son-of-tudor (*sot*) and a line that expressed green fluorescent protein at the *Mst35Ba* and *Mst35Bb* loci (*gfp*). Focal females in our experiment came from the *dah* population, focal males came from the *gfp* population, and competitor males (i.e. the male in the female’s post-mating environment) came from the *sot* population. Males in the *gfp* lines have sperm heads tagged with green fluorescent protein (Manier et al, 2010). *sot* line males produce only seminal fluid, ensuring no sperm transfer in the treatment where females were kept with a competitor male. To generate experimental flies, we used a standard larval density method (following Clancy and Kennington, 2001). Experimental flies were collected within six hours of eclosion to ensure virginity using ice anaesthesia. All flies used in our experiments were between 7-11 days old and kept on a 12:12 light cycle at 25°C, and fed on Lewis medium supplemented with *ad libitum* yeast and molasses.

### Experimental design

Mating assay-We first mated *dah* females to *gfp* males. For this, virgin *gfp* males were introduced individually into vials, each containing a single virgin *dah* female, and their mating behaviours were recorded via continuous scanning of vials. We recorded: mating latency and copulation duration. Once copulation ceased, the mated *dah* female was immediately transferred to a plastic chamber for ejaculate ejection assays (“ejection chamber”).

Ejection assay-The mated females in the ejection chambers were assigned to one of two treatments, either being kept singly in the ejection chamber, or being kept with a virgin *sot* male. Once in the ejection chamber, the females were closely and continuously inspected every 10-15 minutes for up to 12 hours using a dissecting microscope (2x objective magnification) for the presence of an ejected mating plug (Figure 1A, 1B) until ejection occurred. Once an ejected mating plug was observed, it was immediately removed from the mating chamber (Figure 1C) for being imaged the following day (Appendix 1). Additionally, the female was immediately frozen at -20°C to be later dissected and imaged for sperm counts (following Sanghvi et al, 2025, Appendix 2). For all instances of ejaculate ejections, we recorded the ejection latency. One female was observed re-mating with the *sot* male and was therefore excluded from the study. The mating and ejection assays took place over two days (Table S1).

**Figure 1:**
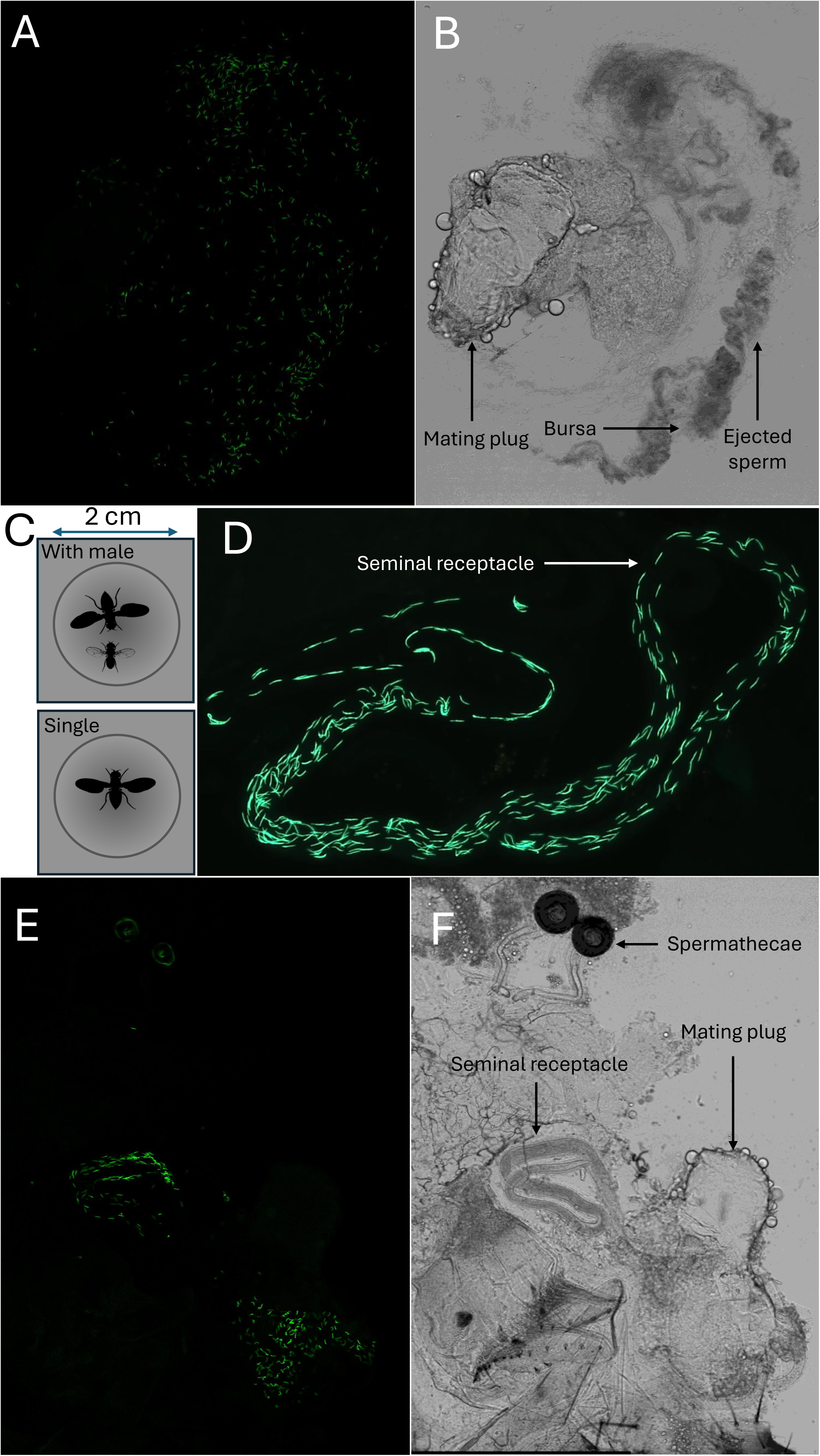
Example of ejaculate ejection and storage, and of ejection chamber treatments. GFP channel (1A) and T-PMT (white light) channel (1B) images of the ejaculate ejected by a female. (1C) Our two experimental treatments showing females either kept singly or with another male inside the ejection chamber. (1D) Sperm inside the seminal receptacle of a female post ejaculate ejection. GFP (1E) and T-PMT (1F) channel images of the reproductive tract of a female who did not eject the ejaculate, thus had mating plug intact and sperm present in the bursa. In 1A, 1C, 1E, green dots/lines show fluorescent sperm heads of individual sperm. Silhouettes taken from PhyloPics are under the <u>CC0 1.0</u> and <u>Public Domain Mark 1.0</u> licences.

Sperm imaging and counts-We dissected the frozen females to count the number of sperm stored in the seminal receptacle post-ejection, and counted the number of sperm in the ejected ejaculate (Appendix 1, 2, Figure 1D-F). Spermathecae sperm counts were omitted due to unclear images, hence only seminal receptacle counts were used as a proxy for sperm retained after ejection. All sperm counts were done using FIJI/ImageJ (Appendix 1, 2). Sample sizes are presented in Supplementary Table S1, STRANGE framework in Appendix 3.

## Data analysis

All analyses were conducted using Rv4.3 (R core team, 2020). We created four models to address our aims. First, we used a Cox-proportional hazards model in the *survival* package (Therneau, 2001) to test how ejection latency (time-to-event dependent variable) and ejection probability (1 or 0 as censoring status) were impacted by the fixed effects of mating latency (aim 1), copulation duration, male presence (aim 2) and day. Copulation duration was included to test whether longer copulations led to more sperm transferred, thus longer ejection latencies. Next, for females that ejected the ejaculate, we constructed two separate Generalised linear models (GLM) with negative binomial error distribution (due to overdispersion in data) using the *glmmTMB* package (Brooks et al, 2017). This model tested how: (i) the numbers of sperm ejected by females (*N_E_*), and (ii) the numbers of sperm retained by females in the seminal receptacle (*N_SR_*), were influenced by the fixed effects of mating latency, copulation duration, male presence, day, and ejection latency. In these models, we included ejection latency to test whether females who ejected the mating plug earlier, ejected more sperm or retained fewer sperm, than females who ejected later (aim 3).

Variation in the numbers of sperm ejected or retained by females could be a consequence of the number of total sperm transferred by the male, rather than merely represent female post-copulatory “choosiness”. In our dataset, sperm numbers in the seminal receptacle were positively correlated with sperm numbers ejected (R^2^ = 0.163, P_lm_ = 0.003, Figure S1), possibly reflecting that females who were inseminated with more sperm, ejected, as well as stored more sperm (Appendix 4). Due to this, we additionally tested how the *proportion* of sperm ejected (*P_E_*) was influenced by our variables of interest, to account for differences in sperm numbers inseminated by males (aim 3). We constructed a GLM with binomial error distribution using the *cbind* function to specify *N_E_* and *N_SR_*, and test how *P_E_* was influenced by the fixed effects of mating latency, copulation duration, ejection latency, day, and male presence. Data in this model were weighted by the total number of sperm counted (i.e. *N_E_* + *N_SR_*), and proportion of sperm ejected was calculated as: 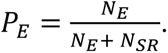

Model assumptions were not violated and were tested using the *DHARMa* (Hartig and Hartig, 2017) *coxme* package (*cox.zph* function).

## Results

### Aim 1

There was no significant influence of mating latency on ejection latency (z = 0.403, P = 0.687, Table S2), *N_E_* (z = -0.418, P = 0.676, Table S3), or *N_SR_* (z = 0.057, P = 0.955, Table S4). However, males with longer mating latencies had smaller proportions of their sperm ejected (*P_E_*) by females, compared to males with shorter mating latencies (z = -2.292, P = 0.022, 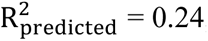, Table S5, Figure 2A).

**Figure 2:**
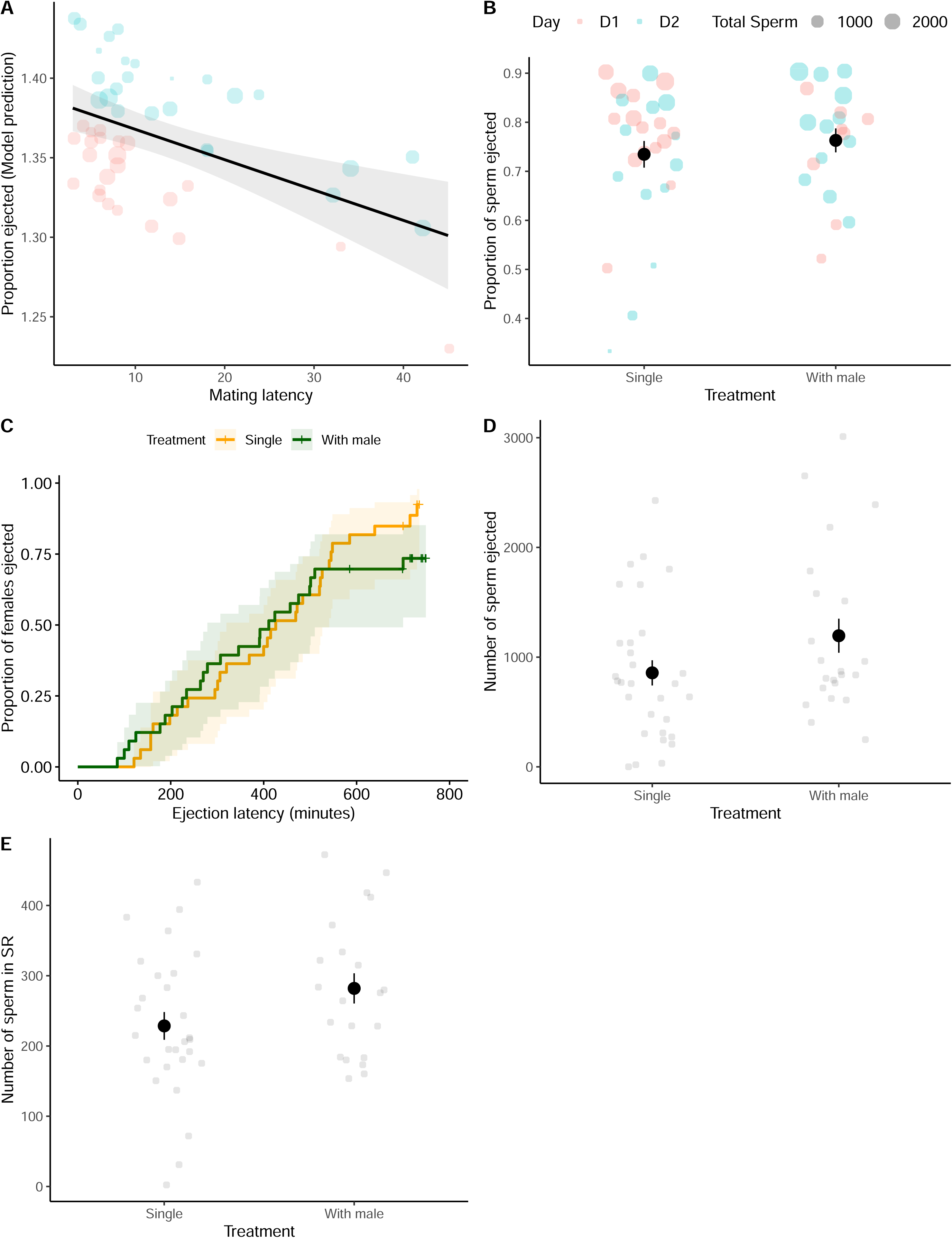
Influence of pre-copulatory variables and male presence in ejection chamber, on post-copulatory female traits. 2A: Longer mating latencies reduced the proportion of sperm (odds ratio) ejected by the female. No significant effect of male presence on 2B: the proportion of sperm ejected; 2C: ejaculate ejection latency; 2D: the numbers of sperm ejected along with the mating plug; 2E: the numbers of sperm stored in the female’s seminal receptacle. Error bars show SEM, shaded areas show 95% C.I. Red dots show data from day 1, blue dots show data from day 2. “+” in 2C shows left-censored individuals.

### Aim 2

Presence of a male in the female’s post-mating environment did not impact: ejection latency (z = -0.689, P = 0.491), *N_E_* (z = 0.981, P = 0.327), *N_SR_* (z = 1.068, P = 0.286), or *P_E_* (z = - 1.345, P = 0.179) (Figure 2B, 2C, 2D, 2E; Table S2-S5).

### Aim 3

Ejaculate ejection latency did not significantly correlate with *N_E_* (z = -1.116, P = 0.264), *N_SR_* (z = -1.367, P = 0.172) or *P_E_* (z = 0.491, P = 0.623).

## Discussion

### Aim 1

We investigated the relationship between pre- and post-copulatory traits related to female choosiness. We found that males who had longer mating latencies also produced sperm that were preferentially stored by females in their seminal receptacle. This result might be due to selection acting on pre- and post-copulatory traits in opposite directions (i.e. disruptively), where males that might have lower success at a pre-copulatory stage, have higher success at the post-copulatory stage (reviewed in Evans and Garcia-Gonzalez, 2016). This could reflect underlying trade-offs between investment in traits that improve mating success or expedite time-to-mating (e.g. courtship-related traits) versus traits that improve fertilisation success and sperm retention by females (e.g. seminal fluid quantity or quality, sperm motility or viability) (Arbuthnott, 2018; Sanghvi et al, 2024). Such underlying trade-offs between pre- and post-copulatory traits could explain why variation in sexually selected traits is maintained despite selection reducing variation (Roff and Fairbairn, 2007). Such negative relationships between pre-versus post-copulatory traits have been shown by Durrant et al (2016) in hissing cockroaches, where males with larger weapons have smaller testes. Our results somewhat contrast those of Doubovetzky et al (2024), who show that fruit fly females are more likely to retain sperm from male strains that they find more attractive pre-copulation, compared to less attractive male strains. This discrepancy might be due to variation in mating latency in our experiment being intra-strain rather than inter-strain.

In fruit flies, male mating latency is frequently used as a proxy for female pre-copulatory choosiness (e.g. Dukas, 2005; Hosken et al, 2008; Kohlmeier et al, 2021; Poissonnier et al, 2024; Robinson et al, 2012; Sanghvi et al, 2025; Savic-Veselinovic et al, 2017; Sharma et al, 2010; Vega-Trejo et al, 2024), while ejaculate ejection latency for post-copulatory choosiness (e.g. Doubovetzky et al, 2024; Yun et al, 2024). However, variation in both mating and ejection latencies can also be a consequence of male-driven processes (Pitnick and Brown, 2000). For instance, males who are more coercive or better at harassing females, might have shorter mating latencies despite being less attractive (Lovlie and Pizzari, 2007). Similarly, due to sperm motility being essential for sperm to enter female storage organs (Holt and Fazeli, 2016; Schnackenberg et al, 2012), males who have more motile sperm might have a better likelihood of their sperm being stored in the seminal receptacle, without requiring any female agency. Thus, the negative correlation between mating latency and *P_E_* might be due to other plausible mechanisms. For example, females could be ejecting more sperm of less-preferred males (who are however, better at coercing females into mating). This is observed in red junglefowl, where subordinate males sneakily coerce females into mating, but have a higher likelihood of having their sperm ejected than sperm of dominant males (Pizzari and Birkhead, 2000). Additionally, mate familiarity confounded with mating latency in our experiment, which could also explain our results.

### Aim 2

We tested how the presence of a male in the female’s post-copulatory environment influenced her ejaculate ejection behaviour. We predicted that due to fruit flies often facing high risk of ejaculate limitation (Sanghvi et al, 2025), or due to females mate-multiplying to increase the genetic diversity of offspring (Jennions and Petrie, 2000), females would eject sperm faster, or eject more sperm, in the presence of a male than in the absence of one. However, we found no significant impact of male presence on any post-copulatory female behaviour. This could be due to females having sufficient sperm from their matings with virgin *gfp* males, therefore not facing any risk of sperm depletion. Doubovetzky et al (2024) recorded ejaculate ejection latency in fruit flies in response to male presence using two different strains. They found that while in one strain, there was no difference in ejection latency between females being kept with males or in isolation, in another strain, females kept with males from the same strain ejected sperm sooner, but when kept with males from another strain, did not. Therefore, our non-significant results for male presence could be due to the specific strains chosen for our experiments, and the strains of males versus females used being different. Yun et al (2024) on the other hand showed that female flies, in the presence of male pheromones in their post-mating environment, eject sperm sooner than flies not exposed to male pheromones. In our study, we manipulated male presence not through odour cues, but directly, which could lead to such discrepancies. In our study, consequently, male presence could have led to male harassment and prevented females from ejecting sperm, thereby cancelling the effect of male [pheromone] presence promoting ejaculate ejection as found by Yun et al (2024).

### Aim 3

We explored how different measures of female ejection behaviour correlate. Studies typically use ejection likelihood (Pizzari and Birkhead, 2000), ejection latency (e.g. Lupold et al, 2013; Manier et al, 2020; Yun et al, 2024), or absolute sperm numbers ejected or stored (e.g. Cordoba-Aguilar, 2006; Doubovetzky et al, 2024) as proxies for female post-copulatory choosiness. However, few have tested whether these measures correlate, and none, to our knowledge, have used the proportion of sperm ejected as a proxy, despite other measures not necessarily reflecting female choosiness. For instance, if sperm from higher quality males reach storage faster, females may eject their sperm sooner simply because storage fills up earlier. Here therefore, ejection latency wouldn’t indicate a preference for lower quality males. Similarly, if larger ejaculates result in more sperm being ejected (our results; also see Dean et al, 2011), then sperm ejection numbers may reflect quantity rather than female preference. Based on our results we suggest that the proportion of sperm ejected is a more biologically meaningful metric of female post-copulatory strategies (Appendix 4). We were unable to measure sperm stored in the spermathecae due to these images being unclear. Studies could replicate our experiment using larger sample sizes, while also comparing sperm numbers inside spermathecae to those in the seminal receptacle, to test whether sperm numbers between these organs differ under varying environments.

## Supplementary material

Appendix (1-4), Supplementary tables (S1-S5) and figures (S1) are provided along with this manuscript in the Supplementary materials.

## Data and code availability

All data from our experiment and the R code used for analysis can be found at OSF: 10.17605/OSF.IO/GDHYQ and https://osf.io/gdhyq/. Images of dissected females’ SR and their ejected ejaculates can be found on Figshare: 10.6084/m9.figshare.28164263; 10.6084/m9.figshare.28164290.v1.

## Author contributions

KS, IS designed the study. KS conducted the experiment, collected and analysed data. BT tested for repeatability of sperm counts. KS wrote the first draft. IS supervised the experiment. IS, BT contributed toward subsequent revisions.

## Acknowledgements

We are grateful to members of the fly lab for helpful suggestions, Ben Hopkins for making the ejection chambers, Umut Calkam for modifying these, Michael Thom for giving *gfp* flies, and Jennifer Holter for helping with confocal microscopy.

## Supplementary material

### Supplementary figures

**Figure S1:**
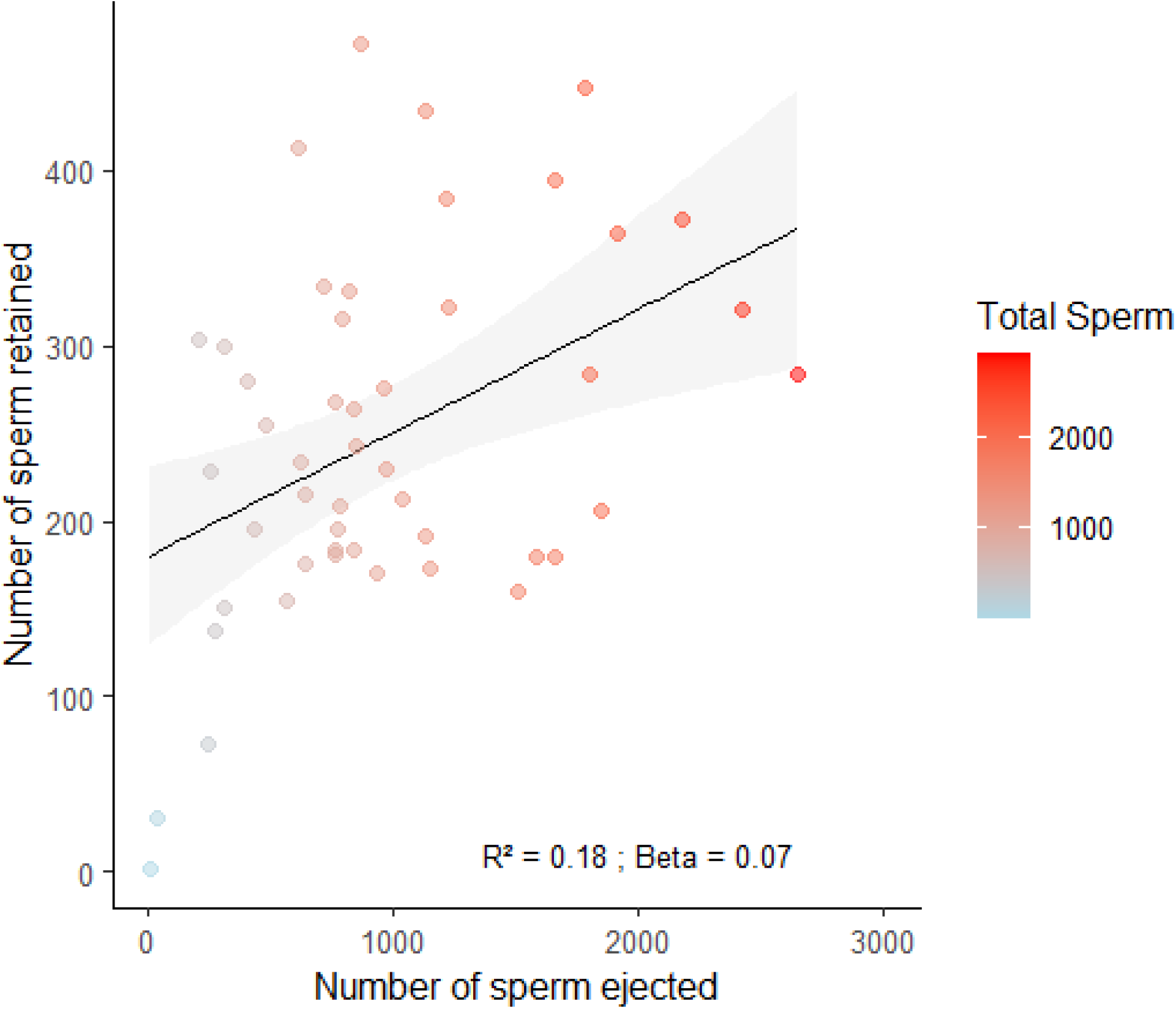
Numbers of sperm ejected by females correlates positively with the number of sperm retained by females, likely due to substantial variation in total sperm numbers transferred by males.

### Supplementary tables

**Table S1:** Sample sizes for different assays in our experiment across the two experimental days and treatments. Mated column represents the number of male-female pairs which mated, thus for whom data on mating latency and copulation duration were obtained. Ejected column represents the number of females that ejected the ejaculate, thus for whom ejection latency data was obtained. Ejected sperm counts column represents the number of females for whom the ejected sperm numbers could be counted. SR sperm counts column represents the number of females for whom sperm numbers in the seminal receptacle were counted. Due to some females/samples being lost at different stages, sample sizes are lower in later assays. On day one, experimental flies were between 7-9 days old and females were transferred into the ejection chambers using an aspirator. One day two, experimental flies were between 9-11 days old and females were transferred into and out of the ejection chamber using carbon dioxide anaesthesia lasting between 5 to 10 seconds. Carbon dioxide was used to reduce the number of females that escaped when using an aspirator (which was the case on day 1).

|  | Mated | Ejected | Ejected sperm counts | SR sperm counts |
| --- | --- | --- | --- | --- |
| **Day 1** |  |  |  |  |
| Single | 17 | 16 | 16 | 15 |
| With male | 14 | 9 | 9 | 8 |
| **Day 2** |  |  |  |  |
| Single | 16 | 14 | 14 | 13 |
| With male | 19 | 15 | 14 | 13 |

**Table S2:**
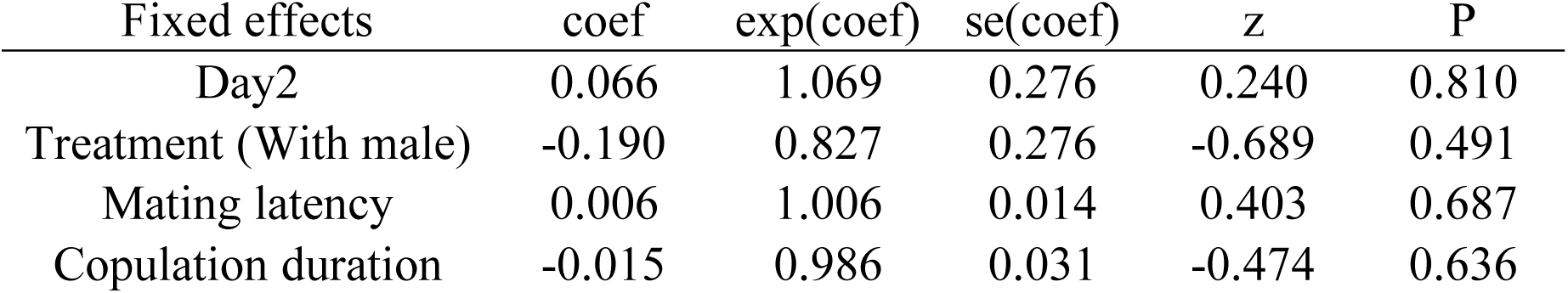
Influence of mating latency (used as a proxy for pre-copulatory choosiness of females), presence of male, copulation duration, and day, on ejection latency. Coz proportional hazards model.

| Fixed effects | coef | exp(coef) | se(coef) | z | P |
| --- | --- | --- | --- | --- | --- |
| Day2 | 0.066 | 1.069 | 0.276 | 0.240 | 0.810 |
| Treatment (With male) | -0.190 | 0.827 | 0.276 | -0.689 | 0.491 |
| Mating latency | 0.006 | 1.006 | 0.014 | 0.403 | 0.687 |
| Copulation duration | -0.015 | 0.986 | 0.031 | -0.474 | 0.636 |

**Table S3:**
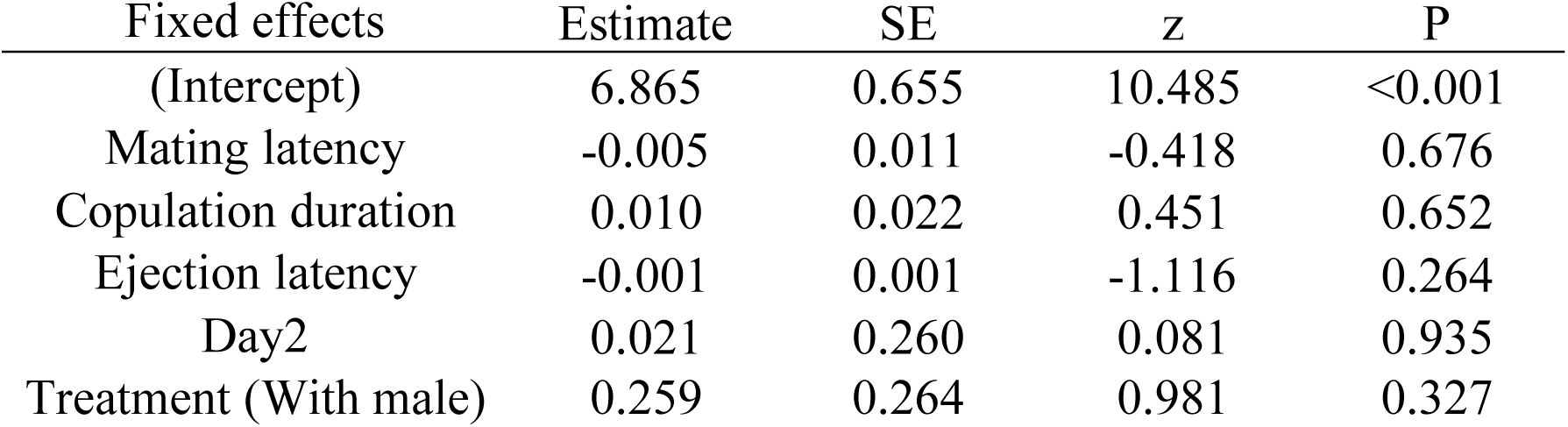
Influence of mating latency, presence of male, copulation duration, day, and ejection latency, on the numbers of sperm ejected (*N_E_*) along with the mating plug. Model constructed as GLM with negative binomial error distribution.

| Fixed effects | Estimate | SE | z | P |
| --- | --- | --- | --- | --- |
| (Intercept) | 6.865 | 0.655 | 10.485 | <0.001 |
| Mating latency | -0.005 | 0.011 | -0.418 | 0.676 |
| Copulation duration | 0.010 | 0.022 | 0.451 | 0.652 |
| Ejection latency | -0.001 | 0.001 | -1.116 | 0.264 |
| Day2 | 0.021 | 0.260 | 0.081 | 0.935 |
| Treatment (With male) | 0.259 | 0.264 | 0.981 | 0.327 |

**Table S4:**
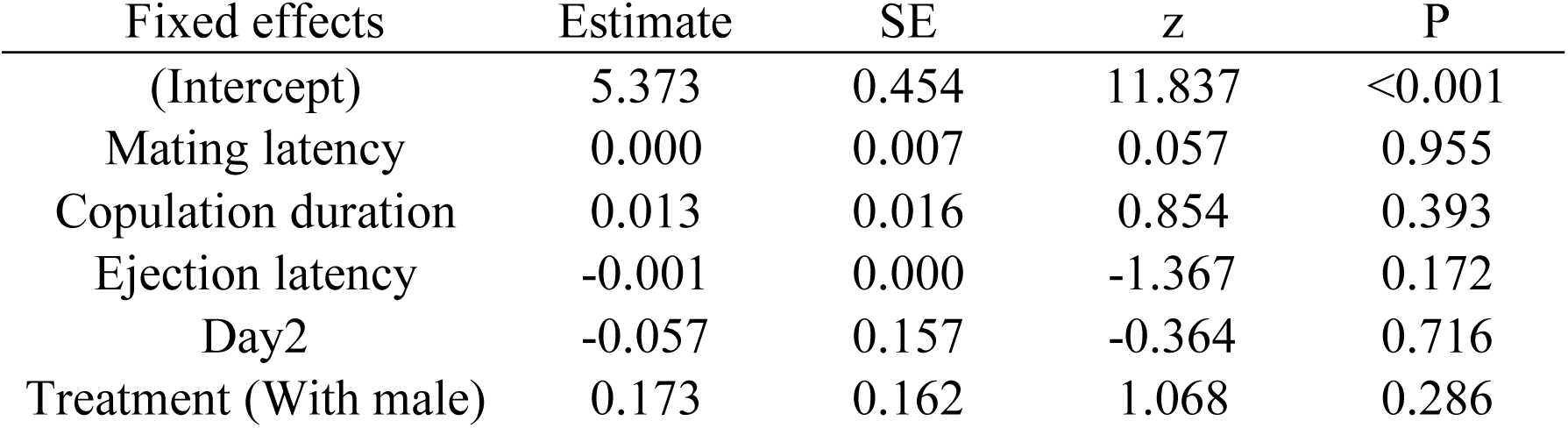
Influence of mating latency, presence of male, copulation duration, day, and ejection latency, on the numbers of sperm stored in the female’s seminal receptacle (*N_SR_*) after her ejecting the mating plug. Model constructed as GLM with negative binomial error distribution.

| Fixed effects | Estimate | SE | z | P |
| --- | --- | --- | --- | --- |
| (Intercept) | 5.373 | 0.454 | 11.837 | <0.001 |
| Mating latency | 0.000 | 0.007 | 0.057 | 0.955 |
| Copulation duration | 0.013 | 0.016 | 0.854 | 0.393 |
| Ejection latency | -0.001 | 0.000 | -1.367 | 0.172 |
| Day2 | -0.057 | 0.157 | -0.364 | 0.716 |
| Treatment (With male) | 0.173 | 0.162 | 1.068 | 0.286 |

**Table S5:** Influence of mating latency, presence of male, copulation duration, day, and ejection latency, on the proportion of ejaculated sperm ejected by the female in her seminal receptacle (*P_E_*) along with the mating plug. Proportion of sperm ejected calculated as numbers retained in seminal receptacle, divided by the sum of numbers retained in seminal receptacle plus numbers ejected: *P_E_* = *N_SR_*/*(N_SR_* + *N_E_*). Model constructed as GLM with binomial error distribution with *N_SR_* + *N_E_* included as weights.

| Fixed effects | Estimate | SE | z | P |
| --- | --- | --- | --- | --- |
| (Intercept) | 1.377 | 0.066 | 20.814 | <0.001 |
| **Mating latency** | **-0.002** | **0.001** | **-2.292** | **0.022** |
| Copulation duration | -0.001 | 0.002 | -0.387 | 0.699 |
| Ejection latency | 0.000 | 0.000 | 0.491 | 0.623 |
| **Day2** | **0.069** | **0.024** | **2.902** | **0.004** |
| Treatment (With male) | -0.032 | 0.024 | -1.345 | 0.179 |

## Appendix

### Appendix 1: Sperm counts in seminal receptacle

Females were dissected using Inox Biology forceps, by pulling apart their last segment from the rest of the body, to separate their reproductive tracts Female reproductive tracts were then placed on a slide coated with an aqueous solution of gelatine and chromium potassium sulphate. The seminal receptacle and spermathecae were spread to allow clear imagining. Then, a coverslip was placed on the sample and the sides of the coverslip were glued using rubber cement to prevent desiccation. The seminal receptacle, bursa, and spermathecae on these slides were imaged on the same day of dissections, using a Nikon Eclipse50i fluorescence microscope (magnification = 10x objective, 10x eyepiece, wavelength = 480nm) with a chromix HD camera, under UV light from a CoolLED pe300 light source. The number of sperm heads (which appeared as fluorescent green under UV light) were later counted manually from images, using the cell counter plugin on FIJI/ImageJ version win32 (Schindelin et al, 2012). Sperm counts were highly repeatable between two different analysts (R² = 0.98 across 11 images). Sperm images of the spermathecae were unclear leading to inaccurate counts, therefore only data on sperm stored in seminal receptacles were used.

### Appendix 2: Sperm counts in ejected mating plug

4µL of PBS was poured on the ejected ejaculate in the chamber, and the mating plug along with the ejected sperm were collected using minutien 0.1mm micropins, and transferred immediately onto a slide coated with an aqueous solution of gelatine and chromium potassium sulphate, and a coverslip was placed on the slide thereafter. The sperm present in the ejected ejaculate were carefully spread-out using microneedles to improve visualisation. The sides of the coverslip were glued using rubber cement to prevent desiccation. On the following day, these sperm were imaged using a confocal Leica stellaris microscope (20x objective, 10x eyepiece magnification) through a GFP channel (excitation wavelength: 480-520nm, gain: 230-260 units, pixel size: 1024×1024, scan speed: 200, pinhole: 1AU), as well with a T-PMT channel (white light). The number of fluorescent sperm heads were later estimated automatically using the find maxima plugin on FIJI/ImageJ version win32 (Schindelin et al, 2012). The prominence on the find maxima plugin was set at 65 because this where the power function relationship between prominence values and estimates of sperm number became linear and flat (following Sanghvi et al, 2025). Sperm counts done manually on images of ejected sperm were highly repeatable with sperm counts estimated automatically using the find maxima plugin (R² = 0.984 based on 16 images), therefore ensuring accuracy of automated sperm counts. Additionally, sperm counts done manually on 16 images of ejected sperm were highly repeatable between two analysts (R² = 0.99).

### Appendix 3: STRANGE framework

Social background-Parents of experimental individuals had been kept in mixed-sex, freely mating, populations cages in a temperature and humidity controlled lab. Experimental males and females were kept as virgins in single-sex vials of 10 individuals, and did not interact with individuals of the opposite sex until the mating assay.

Trappability and self-selection-Experimental flies were reared using a random subset of eggs acquired from the population cages, laid by stock flies in those cages. Post-eclosion, a random subset of flies were collected as virgins for our experiment.

Rearing history-Experimental males only interacted with other virgin experimental males in groups of 10. They were kept in transparent plastic vials with food and without any other objects. Experimental females were kept singly.

Acclimation-Flies were kept in vials for 7 to 11 days before being used for assays.

Natural changes in responsiveness-The lab was 24/7 temperature controlled at 25°C and on a 12:12hr light-dark cycle.

Genetic makeup-Experimental males of *gfp* background had transgenes that expressed green fluorescent protein and had been reared in our lab at least for the past eight years. These flies had been previously backcrossed in *dah* background. For most of this time, *gfp* flies were reared at 20°C, however, for ∼4 generations prior to the experiment, were maintained at 25°C. Females were from outbred *dah* strain and were derived from populations of wild-type flies collected in Benin (Africa) and maintained in our lab since the 1970’s. *sot* males were obtained by crossing two parental lines (Tudor and *dah*) following Sanghvi et al (2025) and only persist for a single generation due to being infertile.

Experience-Experimental flies had not participated in any previous experiments.

### Appendix 4: Assaying sperm ejection

An assumption is that if females are ejecting more sperm, they should have fewer sperm retained, leading to a negative correlation between numbers of sperm ejected versus retained. Therefore, by measuring just one trait (either numbers retained or ejected), studies assess female post-copulatory preference.

We think that this view is incorrect, and suggest that the mathematics developed in life-history theory that considers resource allocation and acquisition (see Roff and Fairbairn, 2007; Van Noordwijk and De Jong, 1986), can be implemented to better understand sperm ejection behaviour. These theories show that the correlations between two life-history traits can be positive or negative, depending on the amount of variation between individuals in a population there is, in resource allocation towards each trait versus resource acquisition. Specifically, if there is more variation in resource allocation (i.e. individuals differ more in how they are partitioning resources to each trait), then the traits themselves will correlate negatively. However, if individuals differ more in how much energy they have, the traits will correlate positively (see Figure S1).

Following these life-history models, sperm ejection behaviour can be represented as:

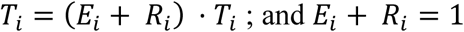

where *T*_*i*_ is the total sperm transferred to the *i*^*th*^ female, *E*_*i*_ and *R*_*i*_ are the *proportions* of inseminated sperm ejected and retained respectively by her; *E*_*i*_ · *T*_*i*_ is the *numbers* of sperm ejected, *R*_*i*_ · *T*_*i*_ is the *numbers* of sperm retained.

Here, if resource acquisition (i.e. the total numbers of sperm inseminated by males to females “T”) varies more between females than resource allocation (i.e. proportion of sperm ejected versus sperm retained, i.e. “*P*_*E*_” or “*P*_*R*_”), then a positive correlation between numbers of sperm ejected versus retained would ensue. The opposite, i.e. more variation in allocation than acquisition would lead to a negative correlation between numbers of sperm retained versus ejected. Thus, measuring either absolute sperm numbers ejected or retained is not informative about the underlying ejection strategy, due to their correlations (positive or negative) being vulnerable to the amount of variation in total sperm inseminated.

For example, when a study only assays the numbers of sperm ejected and finds that females eject fewer sperm of males from treatment A than B, it does not necessarily mean that males from treatment A are preferred. Instead, this result could also be a consequence of males in treatment A inseminating fewer sperm overall. This would lead to males in treatment A also having fewer sperm retained, and here, if another study only assayed the numbers of sperm retained, the opposite conclusion about preference would be reached.

We therefore suggest that the ***proportion*** of total sperm ejected (or retained) should be used to quantify ejection behaviour. This metric accounts for the total numbers of sperm inseminated (i.e. resource acquisition), ejected, as well as retained (resource allocation).

